# Pediatric glioblastoma - unlike normal cells - are sensitive to the combination of vorinostat and olaparib and to its downstream effector - phosphorylated eIF2α

**DOI:** 10.1101/2020.11.25.397497

**Authors:** Karin Eytan, Ziv Versano, Moshe Leitner, Shoshana Paglin, Amos Toren, Michal Yalon

## Abstract

Current therapies offer only a short relief for patients with pediatric glioblastoma (PED-GBM). Therefore, expanding treatment options for this fatal disease is of utmost importance. We found that PED-GBM cell lines, originated from diffuse intrinsic pontine glioma expressing H3K27M mutation (DIPG), or from hemispheric glioma expressing H3G34R mutation, are sensitive to combinations of histone deacetylase and PARP-1 inhibitors (vorinostat with either olaparib or veliparib). These combinations led to an enhanced decrease in their survival, and to increased phosphorylation of eIF2α. Experiments with the S51D phosphomimetic variant of eIF2α and with brain-penetrating inhibitors of phosphorylated eIF2α (p-eIF2α) dephospohrylation, salubrinal and raphin1, showed that increased eIF2α phosphorylation diminished PED-GBM cell survival and sensitized them to PARP-1 inhibitors as well as to ionizing irradiation, which is the main treatment modality in these patients. PED-GBM cells were also remarkably more sensitive to combination of vorinostat and PARP-1 inhibitors and to salubrinal and raphin1 than normal human astrocytes and fibroblasts.

Importantly, although the overall effect of increased eIF2α phosphorylation was a reduced survival of PED-GBM cells, it also increased the cellular level of MTH1, an enzyme that protects treated cells against the incorporation of oxidized nucleotides into nucleic acids, resulting in an enhanced decrease in cell survival in response to the combination of salubrinal and MTH1 inhibitor, TH588.

Our results indicate that combinations of the FDA approved drugs, vorinostat and either veliparib or olaparib, could potentially be included in PED-GBM treatment protocols and that the effect of salubrinal and raphin1 on PED-GBM survival warrants further evaluation.

## Introduction

Glioblastoma is the most severe subtype of human primary brain tumor in both adults and children [1–3]. However, in contrast to adult glioblastomas, a significant percentage of pediatric glioblastomas (PED-GBM) are located in the midline region of the brain and are either completely or partially inaccessible for surgical resection. Further, in pediatric patients, the addition of temozolomide did not show survival benefit compared to radiation therapy alone [4].

The lack of effective treatments for glioblastoma patients prompted a search for new therapies. In particular, a great effort has been directed towards understanding the molecular characteristics of PED-GBM, which could provide clues to its etiology, and elucidate new molecular targets for drug development [5–7]. These studies revealed the existence of somatic heterozygous histone mutations that alter the distribution of histone marks and gene expression, and are mutually exclusive. Thus, the H3.3K27M or H3.1K27M mutations are associated with diffuse midline gliomas (DMG), including diffuse intrinsic pontine glioma (DIPG) and diffuse thalamic gliomas, whereas the H3.3G34R/V/D mutations are harbored by hemispheric gliomas. The mutated histones change the chromatin marks and consequently gene expression [5, 8–10]. The H3K27M mutation in DIPG is associated with overall reduction in the level of H3K27me^3^. However, H3K27me^3^ polycomb repressive complex (PRC) 2 binding domains do exist in these cells as well as bromodomain and extra terminal (BET) domains, which consist of heterotypic H3K27M-K27ac nucleosomes that bind bromodomain proteins [11, 12]. The activity of both domains is required for the survival of DIPG, as the PRC2 binding domains repress differentiation and the BET domains maintain the proliferative capacity of the cells. It has been demonstrated that drugs that interfere with the activity of histone writers or of proteins associated with DIPG chromatin domains, such as BET-associated proteins, affect the survival of the cells [11–14]. Accordingly, the pan-histone deacetylase inhibitors (HDACi), panobinostat, which reverses the suppressive effect of H3.3K27M on H3.3K27 methylation and increases the level of H3.3K27me^3^, also decreases their survival *in vitro* and in xenograft models [14].

Several laboratories, including ours, have demonstrated that HDACi sensitize breast, ovarian, prostate, myeloid leukemia cancer cells as well as glioblastoma to treatment with PARP-1 inhibitors (PARPi), regardless of their capacity for repair of dsDNA break [15–20]. PARP-1 is known for its role in the repair of single strand DNA breaks, and in slowing fork progression during stress, thus preventing fork collapse and allowing for DNA damage repair. Its activity is also involved in the modulation of chromatin methylation [21–23]. Chornenkyy et al. showed that DIPG cells responded poorly to veliparib but were sensitive to olaparib and niraparib [24]. Niraparib and olaparib are reportedly more cytotoxic than veliparib due to their capacity to trap PARP-1 at damaged DNA and create trapped PARP-DNA complexes that are more cytotoxic than unrepaired single-strand breaks [22]. However, unlike vorinostat and veliparib, which penetrate the blood brain barrier olaparib did not show brain penetration in rats, however it has been detected in orthotopic glioblastoma xenografts and in human brain tumor specimens [25].

We previously showed that the combination of vorinostat and veliparib enhanced the death of breast cancer and GBM cell lines, including KNS42, the H3.3G34R expressing PED-GBM cell line, and the U87 cell line, which was derived from an adult GBM [15, 19]. To the best of our knowledge, to date, the effect of HDACi and PARPi combination on DIPG survival has not been reported, either in pre-clinical or clinical trials.

Here, we expanded our experiments to include DIPG cells and tested the effect of vorinostat on their response to veliparib and olaparib. Identifying downstream effectors of anti-cancer treatments can potentially lead to the finding of new molecular target for development of drugs that could preferentially affect cancer cells and thus diminish treatment-associated side effects. Our previous studies showed that a sustained elevation of p-eIF2α mediated the effect of vorinostat on the sensitivity of breast cancer cells to veliparib and enhanced their response to ionizing irradiation. In the experiments reported here, we focused on the role of increased eIF2α phosphorylation in modulating the survival and response to treatment of PED-GBM cell lines. Our findings indicate effective combinations of blood brain barrier penetrating HDACi and PARPi [25–27] as well as drugs that inhibit eIF2α dephosphorylation, which could potentially prove useful in the treatment of PED-GBM.

## Materials and Methods

### Cell lines

The SF188 cell line was derived from the left temporal lobe tumor of an 8-year-old male patient. It carries a focal deletion of neurofibromatosis 1 at 17q11.2 and was kindly provided in 2014 by Dr. Daphne Haas-Kogan (UCSF, San Francisco, CA). This cell line was authenticated in 2017 by short tandem repeat profiling at the Genomic Center Core Facility (Technion, Haifa) and frozen aliquots were prepared and are typically resuscitated and used for eight weeks. SU-DIPG-VI-GFP-LUC (SU-DIPG-VI) cell line derived from pediatric DIPG primary tumor expressing H3.3K27M was a generous gift from Dr. Michelle Monje (Stanford University, Stanford, CA). The KNS-42 cell line, which was derived form a 16-year-old patient glioma and harbors the H3.3G34V mutation, was obtained in 2019 from the Japanese Collection of Research Bioresources Cell Bank (Osaka, Japan). Following a short propagation period, these cell lines were frozen, and aliquots were resuscitated and used for eight weeks. Fibroblasts were obtained from dermal human fibroblasts, per protocol #7044 (approved by the Institutional Review Board at Sheba Medical Center) and normal human astrocytes (HA) were from ScienCell Research Laboratories (Carlsbad, CA). All cells were routinely tested for the presence of mycoplasma.

### Growth Condition

KNS-42 cells were grown in Eagle’s minimal essential medium, and SF188 cells were grown in high-glucose DMEM. The media were supplemented with fetal bovine serum (FBS) (5% for KNS-42 and 10% for SF188), 1% L-glutamine (KNS-42), penicillin, and streptomycin (Biological Industries, Kibbutz Beit-Haemek, Israel). SU-DIPG-VI cells were grown in tumor stem medium (TSM) consisted of 50% Neurobasal-A Medium, 50% DMEM/F-12, 1% HEPES (1M), 1% sodium pyruvate, 1% non-essential amino-acids, 1% GlutaMAX and 1% antibiotic antimycotic, 2% B-27 (Thermo Fisher Scientific, Waltham, MA), 0.02% heparin (STEMCELL Technologies, Canada), 20 ng/ml EGF and bFGF, and 10 ng/ml PDGF-AA and PDGF-BB (PeproTech Asia, Rehovot, Israel). HA cells were grown in complete astrocytes medium supplemented with 1% penicillin and streptomycin, 2% FBS, and 1% astrocytes growth supplements (AGS) (all from ScienCell). Fibroblasts were grown in high-glucose DMEM that was supplemented with 20% FBS, penicillin-streptomycin, 1% non-essential amino acids, and 0.2% β-mercaptoethanol (Invitrogen, Life Technologies, Grand Island, NY).

### Reagents

Vorinostat was obtained from LC laboratories (Boston, MA), olaparib was from MedChem Express (NJ), Salubrinal and the MTH1 inhibitor, TH588 (TH), were from Sigma-Aldrich (Merck KGaA, Darmstadt, Germany), raphin1 was from Tocris Bioscience (Bristol, UK). Veliparib was from APExBIO (Houston, TX). All drugs were added from stock solutions in dimethyl sulfoxide (DMSO) and control cultures received an equal amount of the vehicle. The final concentration of DMSO in the culture medium did not exceed 0.1%. All other materials were of analytical grade.

### Survival assays

SU-DIPG-VI cells were plated in triplicates at a density of 3500 cells per well in 96-well plates. Cells from one triplicate were dissociated 24 h later and counted to determine cell number at the time of treatment initiation (T_0_). Eight days later, the number of surviving cells was determined by counting trypan blue excluding cells (Te). Cell survival was expressed in percent according to the following formula:

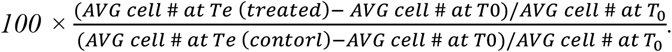

Cell death was calculated as follows: *100 × ((AVG cell # at T0 - AVG cell # at Te)/AVG cell # at T0)).*

The growth rate of astrocytes was higher than that of DIPG. Therefore, to compare the effect of the various treatments on the number of cell divisions in SU-DIPG-VI cells and in HA, we plated both cell types at an equal density of 1200 cells per well, which is lower than the regular density of 3500 cells/well. The cells were plated in triplicates and the number of cells at T_0_ was determine as described above. Trypan blue excluding cells were counted 7 days later (Te) and the inhibition of cell division was calculated according to the following formula:

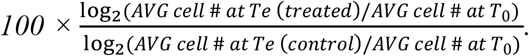

Interestingly, the lower density seemed to increase the sensitivity of DIPG cells to treatments, as noted in comparing the results in Fig. 1–3 to those demonstrated in Fig. 5.

**Fig. 1.**
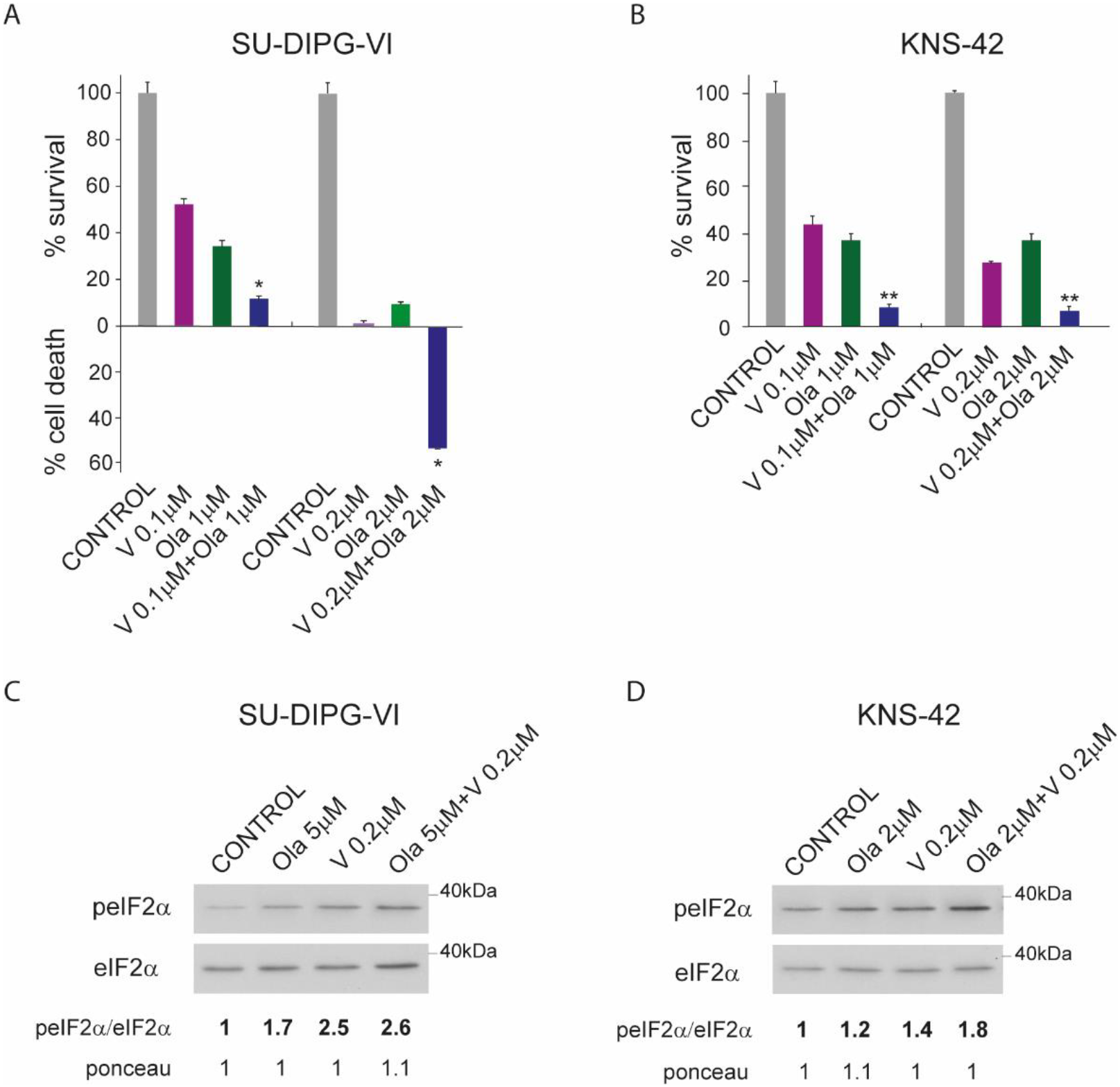
Combined treatment of vorinostat and olaparib leads to enhanced decrease in PED-GBM survival: **A**. Cells (3500/well) were plated in triplicates, treated, incubated and counted as described in Methods. Values are mean survival (%) or cell death (%) ± S.D. Differences between survival of combined treatment and each one of the sole treatments or the control were significant - *p<0.05. **B**. Cells (500/well) were plated in triplicates for clonogenic survival assay and treated as described in methods. Values are mean survival fraction (%) ± S.D. Differences between survival of combined treatment and each one of the sole treatments or the control were significant - **p<0.005. **C**, **D**. Cells were treated and processed for western blot analysis of changes in the ratio peIF2α/eIF2α as described in methods. Numbers at the bottom of the autoradiograms indicate changes relative to controls of p-eIF2α/ eIF2α and of loaded proteins (Ponceau). V-vorinostat, Ola-Olaparib, eIF2α - total eIF2α, peIF2α-phosphorylated form of eIF2α. The experiments were reproduced once with similar results.

Experimental combination indices (CI) were obtained by employing the computer program CompuSyn, according to a method by Chou [28]. CI < 1 indicated synergistic interaction, and C = 1 indicated an additive one. Surviving fraction was defined as the number of treated cells at the end of the experiment (Te) divided by the number of untreated cells (control) at that time.

SU-DIPG-VI images were captured with EVOS^®^FL imaging system (Thermo Fisher Scientific, Waltham, MA) and a 10x objective.

### Clonogenic survival assay

Clonogenic survival assays were conducted in 6-well plates. Cells (500/well) were plated in triplicate. Treatment was initiated 24 h post-plating. When 90-95% of the colonies in the control plate possessed more than 50 cells (~14 days post-plating), colonies were fixed with 70% ethanol and stained with 0.05% crystal violet (Merck) before counting.

### Radiation

Cells were irradiated in an X-ray irradiator (Polaris sc-500 series II) at a dose-rate of 100 cGy/minute.

### Cell lysis and Western blotting

Treatments (addition of drugs or irradiation) were initiated 48 hours post-plating and incubation was continued for the specified time before harvesting cells. Cells were washed with ice-cold DPBS and collected in a buffer containing 150 mM NaCl, 50 mM Tris (pH 7.5), 2% SDS, 1.7 mM EDTA, 1.5 mM sodium orthovanadate, 100mM sodium fluoride, 1.5μg/mL pepstatin, and Roche anti-protease cocktail. Following heating at 95°C and clearing by centrifugation, protein content was determined with a bicinchoninic acid reagent (Bio-Rad, Hercules, CA). Equal loading was verified in a twin run of electrophoresed individual strips of protein stained with Ponceau S (Sigma, St-Louis, MO) and extracted with PBS supplemented with 1.125 M NaCl and 0.6% Tween-20 (Merck) by measuring absorbance at 520 nm [29]. Gels were blotted onto PVDF or nitrocellulose membranes using the Trans-Blot Turbo system (Bio-Rad). Rabbit anti-p-eIF2α, anti-eIF2α, and anti-MTH1 were from Cell Signaling Technology (Danvers, MA) and HRP-coupled goat anti-rabbit IgG was obtained from Jackson Immunoresearch Laboratories (West Grove, PA). Blots were exposed to X-ray film for chemiluminescence, following treatment with a West Pico ECL reagent (Thermo Fisher Scientific). Values for the integrated light density of autoradiograms were obtained with ImageJ software and were employed for the determination of treatment-induced changes in protein levels. All experiments were reproduced at least once.

### Transfection of cells with plasmids coding for eIF2α variants

S51A and S51D heIF2α variant in pcDNA3.CD2 expression vectors were obtained from Addgene (Cambridge, MA). Transient transfection of KNS-42 was conducted 24 h post-plating, and of SU-DIPG-VI at the time of plating, with 0.5 μg plasmids and JetPEI (Polyplus, New York, NY) according to manufacturer’s instruction.

### RNA extraction and Real-time RT-qPCR

RNA was isolated using Direct-zol RNA Miniprep (Zymo research, Irvine, CA). RT-PCR was carried out with 1 μg RNA using the High Capacity cDNA Reverse Transcription Kit (Applied Biosystems, Foster city, CA), according to the manufacturer’s instructions. Real-time RT-qPCR was performed using 2xqPCRBIO SyGreen Blue Mix (PCR Biosystems, London, UK), according to manufacturer’s instructions, in a StepOnePlus™ system (Applied Biosystems) using the primers:

MTH1
F: 5’-CCGTGGAGAGCGACGAAAT-3’
R: 5’-TTCCAGATGAAACCCGGCTC-3’
β-Actin
F: 5’-CCTGGCACCCAGCACAAT-3’
R: 5’-GCCGATCCACACGGAGTACT-3’

Evaluation of treatment-induced changes in RNA level was calculated according to the ΔC_T_ method using β-actin as a reference gene.

### Statistical analysis

The significance of the differences between the surviving fraction of a combination treatment and each one of its components or the untreated control, was verified by employing the unpaired Student’s t test. P < 0.05 in each comparison was considered statistically significant.

## Results

### The combination of vorinostat and PARP-1 inhibitors is deleterious to the survival of PED-GBM

As shown in Figs.1 A, 5 B and S1A, the combination of vorinostat with either olaparib or veliparib led to enhanced decrease in the survival of DIPG cells. The interactions between vorinostat and olaparib was synergistic as manifested by the relevant CIs (Fig. S2A). Similarly, KNS-42 cells responded to the combination of vorinostat and olaparib by enhanced clonogenic death (Fig.1B; Fig. S1C), as they did to the combination of vorinostat and veliparib [19].

### The effect of increased phosphorylation of eIF2α on the survival of PED-GBM

We have previously shown that increased phosphorylation of this initiation factor is detrimental to the survival of breast cancer cells and increases their sensitivity to ionizing irradiation and veliparib [15, 29]. As noted in Fig. 1 (C, D), the combined effect of vorinostat and olaparib on cell survival was also associated with increased eIF2α phosphorylation in PED-GBM cells. To determine the role played by increased eIF2α phosphorylation in modulating cellular response to the combination of vorinostat and olaparib, we monitored cell survival in response to salubrinal and raphin1 (inhibitors of eIF2α dephosphorylation) as well as to the phosphomimetic variant, eIF2α S51D. Both salubrinal and raphin1 led to a dose-dependent increase in p-eIF2α (Fig. 2A, D) and to a decrease in cell survival (Fig. 2B, C, E, F; Fig. S1B, D). A transient transfection with the phosphomimetic (S51D) and the non-phosphorylatable (S51A) eIF2α variants also demonstrated the negative effect of eIF2α phosphorylation on cell survival (Fig. 2G, H). Salubrinal or raphin1 in combination with olaparib led to an increased level of p-eIF2α and to an enhanced decrease in cell survival (Fig. 3A–D). As with vorinostat and olaparib, the interaction between salubrinal and olaparib was synergistic (Fig. S2B). These experiments show that increased eIF2α phosphorylation, in and of itself, decreases the survival of PED-GBM, and participates in modulating the response of these cells to olaparib, as well as to the combination of vorinostat and olaparib.

**Fig. 2:**
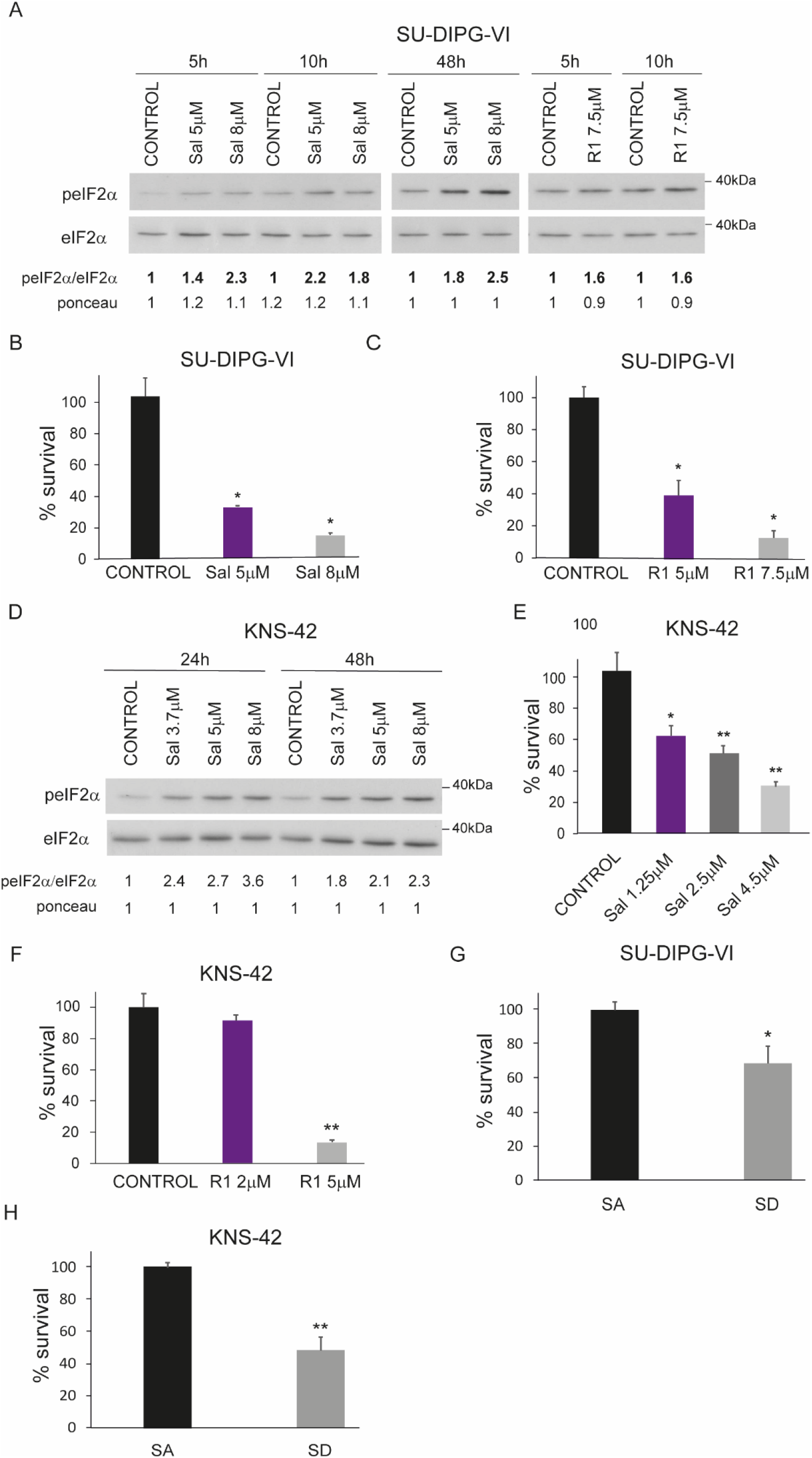
Increased phosphorylation of eIF2α is detrimental to cell survival: *Inhibitors of eIF2α dephosphorylation led to a dose-dependent increase in eIF2α-P*: **A, D**. Cells were treated and processed for western blot analysis of changes in the ratio peIF2α/eIF2α as described in methods. Numbers at the bottom of the autoradiograms indicate changes relative to controls of p-eIF2α/ eIF2α and of loaded proteins (Ponceau). Sal-salubrinal R1 – raphin1. *Increased eIF2α-P decreases cell survival*: *Inhibitors of eIF2α dephosphorylation decrease survival*: **B**, **C, G**. Cells (3500/well) were plated in triplicates, treated with drugs or transiently transfected with eIF2αS51A (SA) or with eIF2αS51D (SD), incubated and counted as described in Methods. Values are mean survival (%) or cell death (%) ± S.D. Differences between survival of combined treatment and each one of the sole treatments or the control were significant - *p<0.05. **E, F, H**. Cells (500/well) were plated in triplicates for clonogenic survival assay and treated with drugs or transiently transfected with SA or with S51D, as described in methods. Values are mean survival fraction (%) ± S.D. Differences between survival of combined treatment and each one of the sole treatments or the control were significant - *p<0.05, **p<0.005. The experiments were reproduced once with similar results.

**Fig. 3.**
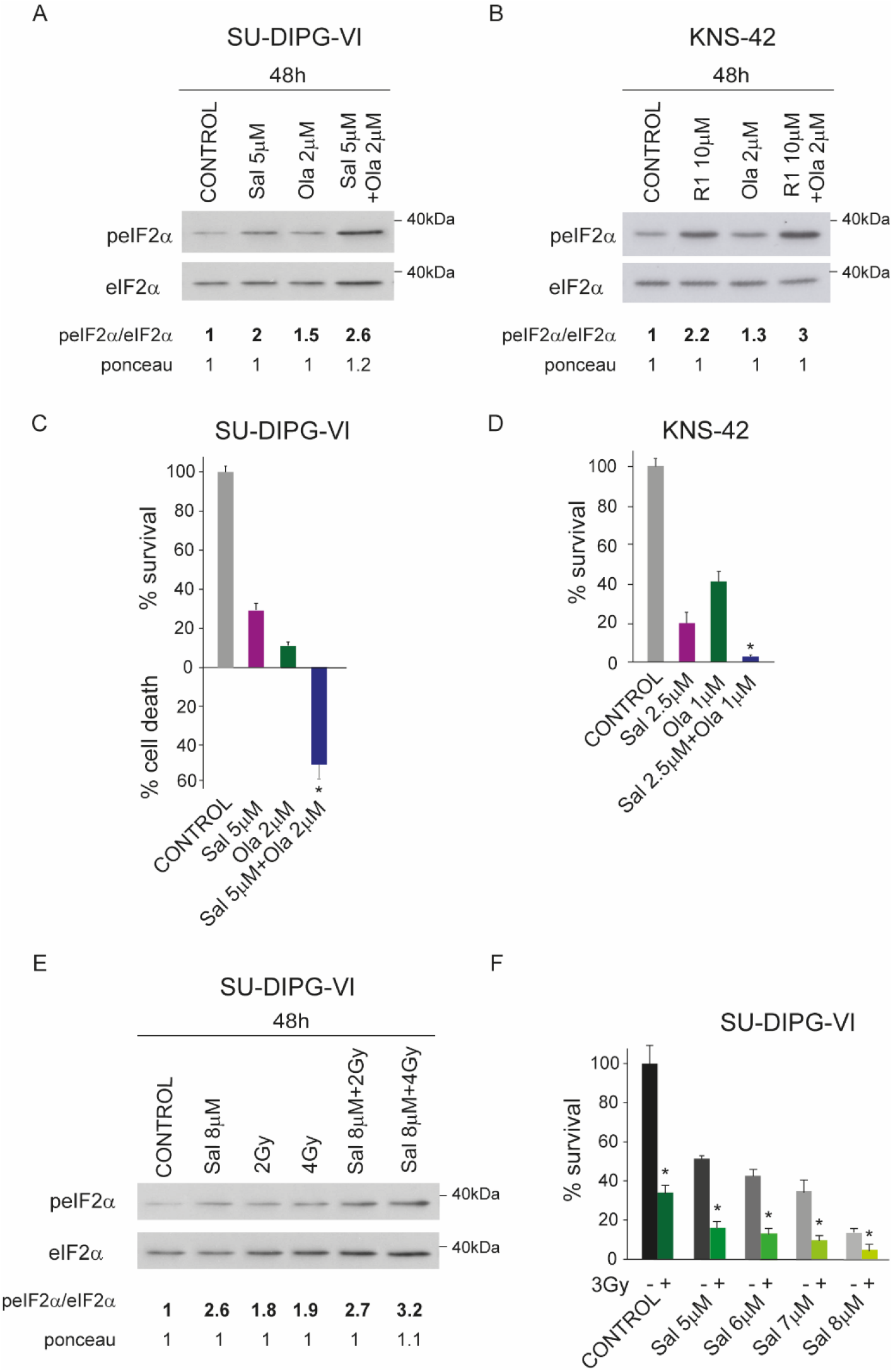
Increased eIF2alpha-P enhances response to Olaparib and radiation in PED-GBM: *Combined treatment of olaparib and salubrinal or raphin1 increases the level of eIF2α-P*: **A**,**B**. Cells were treated and processed for western blot analysis of changes in the ratio peIF2α/eIF2α as described in methods. Numbers at the bottom of the autoradiograms indicate changes relative to controls of p-eIF2α/ eIF2α and of loaded proteins (Ponceau). *Combined treatment of olaparib and salubrinal leads to enhanced decrease in cell survival* **C**. Cells (3500/well) were plated in triplicates, treated with drugs as described in Methods. Values are mean survival (%) or cell death (%) ± S.D. Differences between survival of combined treatment and each one of the sole treatments or the control were significant - *p<0.05. **D**. Cells (500/well) were plated in triplicates for clonogenic survival assay and treated with drugs as described in methods. Values are mean survival fraction (%) ± S.D. Differences between survival of combined treatment and each one of the sole treatments or the control were significant - *p<0.05. **E**. *Combined treatment of radiaiton and salubrinal increases the level of eIF2α-P*: Cells were irradiated and processed for western blot analysis of changes in the ratio peIF2α/eIF2α as described in methods. Numbers at the bottom of the autoradiograms indicate changes relative to controls of p-eIF2α/ eIF2α and of loaded proteins (Ponceau). **F**. *Combined treatment of radiation and salubrinal leads to enhanced decrease in cell survival:* Cells (3500/well) were plated in triplicate, treated with drugs as described in Methods. Values are mean survival (%) or cell death (%) ± S.D. Differences between survival of combined treatment and each one of the sole treatments or the control were significant - *p<0.05. Sal - salubrinal, R1- raphin1, Ola - olaparib, Gy - Gray. The experiments were reproduced once with similar results.

Of importance is the effect of salubrinal on the sensitivity of PED-GBM to ionizing radiation, which is a main treatment modality in DIPG patients. The interaction between salubrinal and radiation was additive, and the combined treatment led to an increased level of p-eIF2α as well (Fig. 3E, F; Fig. S2C).

### Increased phosphorylation of eIF2α leads to increased cellular level of MTH1

The MTH1 enzyme protects treated cells against the incorporation of oxidized nucleotides into nucleic acids thereby playing a role in their resistance to therapy [19, 30, 31]. As depicted in Fig. 4 (A-C, E), increased eIF2α phosphorylation led to increased expression of MTH1. In line with this result, the combination of salubrinal and TH588, an MTH1 inhibitor [32], interacted in a synergistic manner to decrease the survival of PED-GBM (Fig.4D, F; Fig. S2D).

**Fig. 4:**
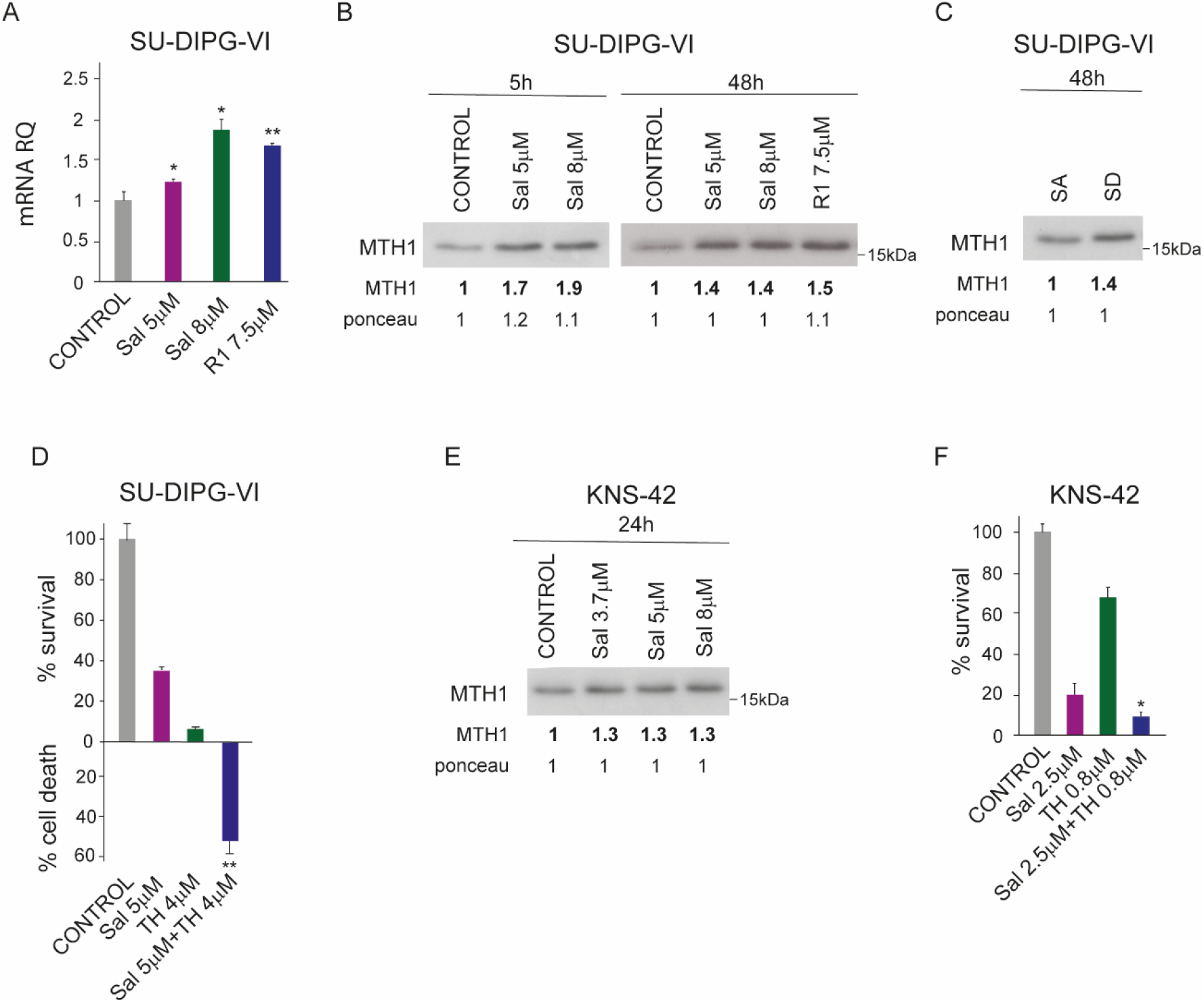
Increased phosphorylation of eIF2α leads to increased level of cellular MTH1. **A.** Cells were treated with salubrinal and raphin 1 for 3.5 hours. RNA was extracted and treatment-induced change in MTH1 mRNA was evaluated by and qRT-PCR as described in Methods. Data are mean relative quantification ± S.D of three triplicates. Differences between treatments and control were significant - *p<0.05, **p<0.005. **B, C, E.** Cells were treated and processed for western blot analysis of treatment-induced changes in cellular level of MTH1 as described in methods. Numbers at the bottom of the autoradiograms indicate changes relative to controls of MTH1 and of loaded proteins (Ponceau). **D.** Cells (3500/well) were plated in triplicates, treated with drugs as described in Methods. Values are mean survival (%) or cell death (%) ± S.D. Differences between survival of combined treatment and each one of the sole treatments or the control were significant - **p<0.005. **F.** Cells (500/well) were plated in triplicates for clonogenic survival assay and treated with drugs, as described in methods. Values are mean survival fraction (%) ± S.D. Differences between survival of combined treatment and each one of the sole treatments or the control were significant - *p<0.05. Sal - salubrinal, R1 - raphin1, SA - eIF2αS51A, SD - eIF2αS51D, TH - TH588. The experiments were reproduced twice with similar results.

### PED-GBM shows higher sensitivity to the combination of vorinostat and olaparib and to increased eIF2α phosphorylation than normal human astrocytes and fibroblasts

As we previously reported, the combination of vorinostat and veliparib [19] or salubrinal barely affects the survival of normal human skin fibroblasts Fig. S1D. As noted in Fig. 5 (A, B), the effect of the combination of vorinsotat and olaparib or veliparib on cell division was much higher in DIPG cells than in astrocytes. The effect of salubrinal and raphin1 on cell division was dramatically higher in DIPG cells than in astrocytes (Fig. 5C, D). In fact, raphin1 did not lead to any significant inhibition of astrocytes cell division. Similarly, the combination of salubrinal and olaparib inhibited cell division in DIPG cells, but had only a minimal effect on astrocytes (Fig. 5E).

**Fig. 5:**
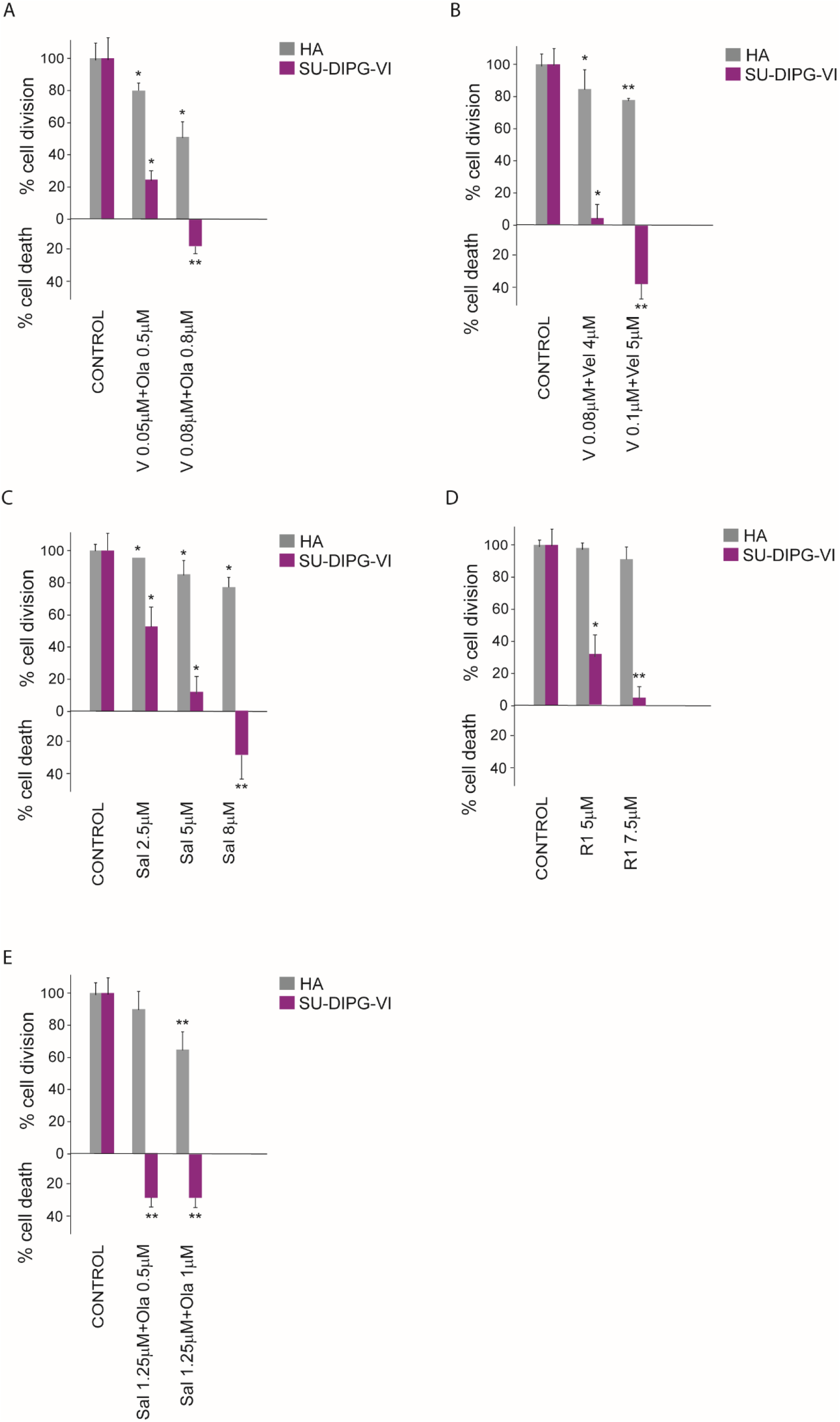
PED-GBM show higher sensitivity to the combination vorinostat and PARP-1 inhibitor and to increased eIF2α-P than normal human astrocytes: Cells (1200/well) were plated in triplicates, treated with drugs as described in Methods. Values are mean number of cell division (%) or cell death (%) ± S.D relative to control untreated cells. Differences between treated and untreated cells were significant - *p<0.05, **p<0.005. The experiments were reproduced once with similar results. HA-human astrocytes, V-vorinostat, Ola – olaparib, Vel - veliparib, Sal-salubrinal, R1 - raphin1.

## Discussion

Our results show that combinations of FDA approved HDACi and PARPi, as well as drugs that inhibit dephosphorylation of eIF2α, decrease survival of PED-GBM with only minimal effect on normal cells. These treatments could potentially expand the available options for the treatment of PED-GBM patients and should be further tested in animal models for their possible use in clinical protocols.

The interaction between vorinostat and PARP-1 inhibitor was synergistic and led to an increased phosphorylation of eIF2α. This observation is similar to that obtained with obtusaquinone, which decreased the survival of DIPG cells both *in vitro* and in orthotopic xenograft model, with a consequent increase in ER stress and eIF2α phosphorylation in these cells [33].

Increased eIF2α phosphorylation is mediated by kinases that respond to various stress signals such as ER overload, viral infection, heme deficiency, and amino acid starvation. It attenuates global protein translation, but simultaneously increases translation of specific mRNAs with either an internal ribosome entry-site element or short open reading frames in their 5’ UTR. These mRNAs code for proteins that modulate the response of the cells to the inflicted stress. It has been suggested that the final outcome of increased eIF2α phosphorylation—whether protective or damaging — is dependent upon the type of stress signal, the kinase that relays it, the duration of the signal, and the cell type [34]. Dephosphorylation of phosphorylated eIF2α is executed by the catalytic PP1 subunit in a complex with G-actin and with either one of two regulatory subunits: the constitutive CReP or the inducible GADD34 [35]. Inhibition of eIF2α dephosphorylation, which leads to an increased p-eIF2α, has been applied in cellular models to assess the role of p-eIF2α in modulating the cellular response to various treatments and metabolic conditions. Increased survival in response to increased eIF2α phosphorylation was considered an evidence that the latter protects the cells against the inflicted damage, whereas increased cell death suggested that it plays a role in exacerbating the inflicted damage [15, 29, 36–39].

Salubrinal was the first inhibitor of eIF2α dephosphorylation to be discovered. The experiments suggested that salubrinal inhibits eIF2α dephosphorylation by interfering with the interaction between PP1 and its regulatory subunits CReP and GADD34 [36]. Recently, a new compound, raphin1 was described. Raphin1 increases eIF2α phosphorylation in cells, and its specific interaction with purified PP1-CReP holophosphatase was demonstrated *in vitro* by employing a surface plasmon resonance-based method [40]. In our studies, we employed salubrinal and raphin1 to evaluate the role played by increased eIF2α phosphorylation in modulating the response of PED-GBM to the combination of vorinostat and PARP-1 inhibitors. We found that both salubrinal and raphin1 increased eIF2α phosphorylation and decreased survival in a dose-dependent manner. We also employed the phosphomimetic S51D eIF2α variants and the non-phosphorylatable S51A variant and demonstrated that the S51D variant decreased cell survival. Importantly, salubrinal and raphin1 increased the sensitivity of PED-GBM to olaparib and to ionizing irradiation.

The eIF2α inhibitors penetrate the blood brain barrier and salubrinal reportedly imparts neuroprotection in rodent animal models following traumatic brain injury or ischemia [41]. However, whereas salubrinal affects memory [42], chronic administration of raphin1 at various doses did not impair the memory of mice in Morris water maze or in a fear-conditioning paradigm [40]. The fact that raphin1, salubrinal, and the combined treatment of salubrinal and olaparib or vorinostat and olaparib lead to a much higher response in DIPG than in astrocytes suggests that evaluation of their effect of on DIPG survival and elucidation of their downstream effectors warrant further studies.

Lastly, in spite of its overall deleterious effect on the survival of PED-GBM, increased eIF2α phosphorylation elevated the cellular level of the MTH1 mRNA and MTH1 protein. Inhibition of this protective enzyme in salubrinal-treated cells led to a synergistic effect on PED-GBM survival. This result is consistent with our previous studies which showed that addition of MTH1 inhibitor to the combination of HDAC and PARP-1 inhibitors enhanced the incorporation of oxidized nucleotides into DNA and cell death [19]. To the best of our knowledge, MTH1 is a newly identified downstream effector of p-eIF2α. This suggests that the effect of MTH1 activity should be taken into account in future studies designed to examine the effect of increased eIF2α phosphorylation on PED-GBM survival.

## Author Contributions

K.E conducted and designed experiments, analyzed data and participated in the writing of the manuscript. Z.V conducted and designed experiments. M.L designed PCR experiments. S.P, A.T and M.Y designed experiments, wrote the manuscript and supervised all aspect of the work.

