## Supplementary material for "Pediatric glioblastoma - unlike normal cells - are sensitive to the combination of vorinostat and olaparib and to its downstream effector - phosphorylated eIF2α": Figure S1 and Figure S2

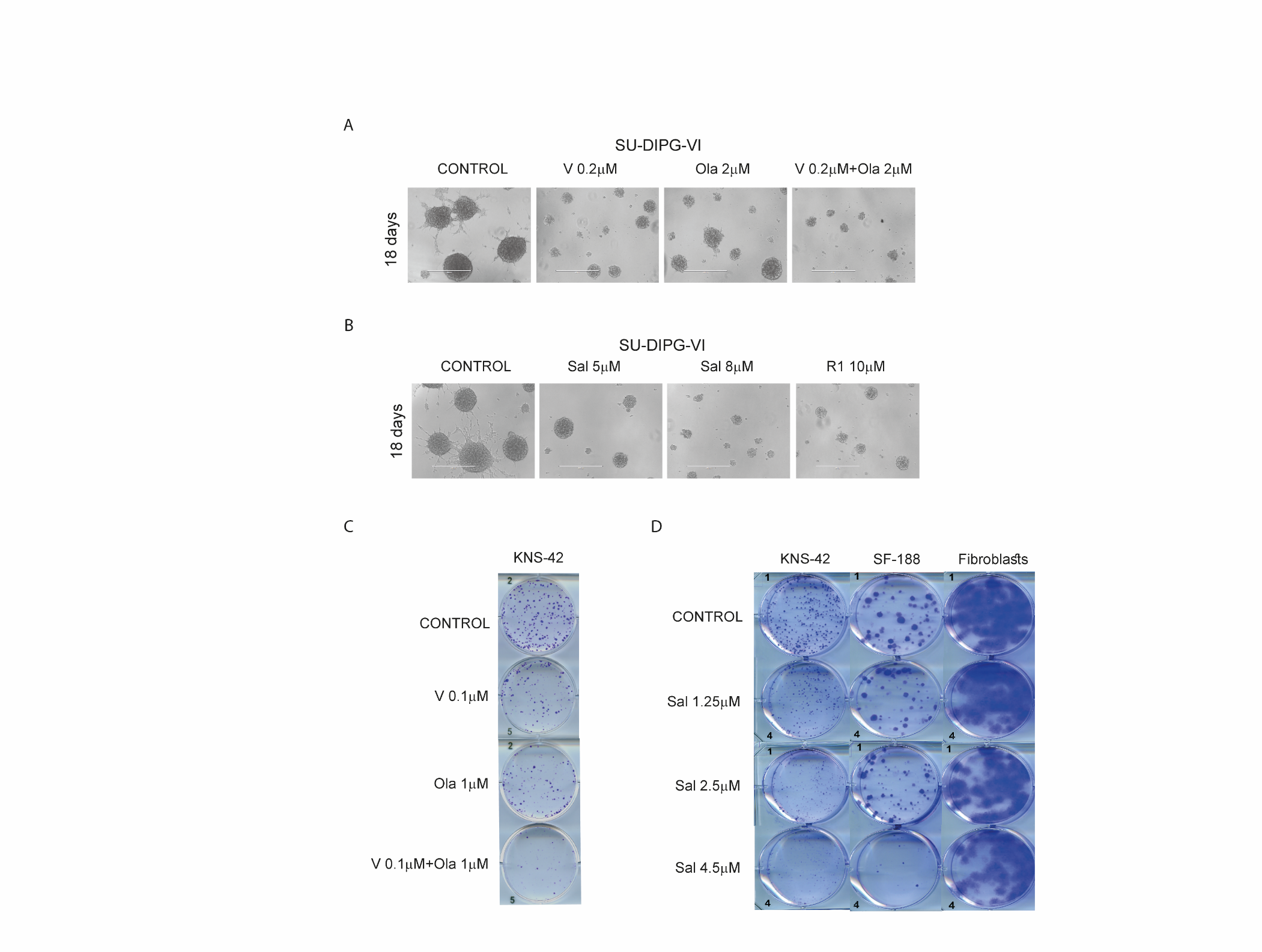

Fig. S1: **Representative images of control and surviving cells**: Control and treated SU-DIPG-VI were exposed to vorinostat, olaparib and their combination (**A**) or to salubrinal and raphin1 (**B**) for18 days post-plating and images were captured as described in Methods. Bar represents 400µm. KNS**-**42, SF188 and normal human skin fibroblasts were plated in triplicates for colony surviving assay, treated with vorinostat, olaparib and their combination **(C),** or with salubrinal **(D)** as described in Methods**.**

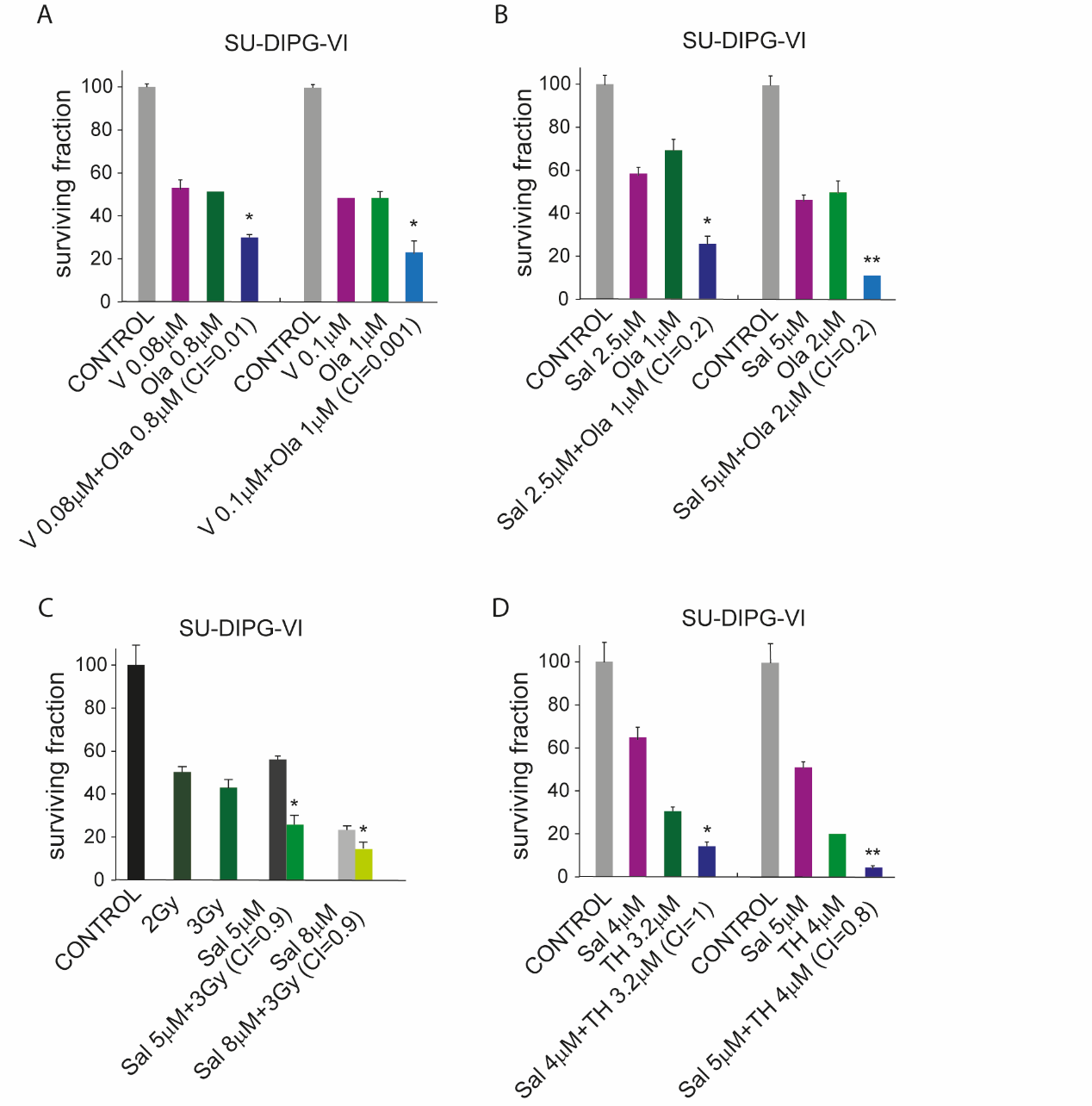

Fig. S2: **Determining the type of interactions between drugs in the various combinations used in this study**: Cells (3500/well) were plated in triplicates, treated with drugs as described in Methods. Values are mean surviving fraction (%) ± S.D. Differences between survival of combined treatment and each one of the sole treatments or the control were significant - *p<0.05, **p<0.05. CI were determined as described in Methods. V-vorinostat, Ola - olaparib Sal- salubrinal, Gy - Gray, TH-TH588
